# Complete mitochondrial genomes do not distinguish phenotypically distinct lineages of Andean *Coeligena* hummingbirds

**DOI:** 10.1101/2020.04.14.041723

**Authors:** Catalina Palacios, Leonardo Campagna, Juan Luis Parra, Carlos Daniel Cadena

## Abstract

Lack of divergence in mitochondrial DNA between species with clear phenotypic differences may be the result of low resolution of markers, incomplete lineage sorting, introgression, or the interplay of various evolutionary mechanisms acting on different traits and genomic regions through time. Previous work revealed that the Andean hummingbirds *Coeligena bonapartei* and *C. helianthea* lack genetic divergence in the mitochondrial *ND2* gene, which shows variation discordant with coloration phenotype but consistent with geography. We sequenced and analyzed complete mitochondrial genomes for *C. b. bonapartei, C. b. consita, C. h. helianthea* and *C. h. tamai* to assess whether patterns revealed by *ND2* analyses hold when considering the entire mitogenome, and to shed light into the evolutionary history of these hummingbirds. We found very low genetic differentiation in mitogenomes among the four lineages of *Coeligena*, confirming patterns based on *ND2* data. Estimates of genetic differentiation, phylogenies and haplotype network analyses of complete mitogenomes did not separate phenotypically distinct taxa, but were consistent with a previously described pattern of northern vs. southern divergence along the Cordillera Oriental of Colombia. Mitogenomes of *C. b. bonapartei* and *C. h. helianthea* are indistinguishable, suggesting incomplete lineage sorting or strong introgression. Mitogenomes of *C. b. consita* and *C. h. tamai* are slightly differentiated, but they are more similar to each other than either is to that of its respective nominate subspecies, a result also suggestive of mtDNA introgression despite distinct phenotypic differences. Our results indicate that various evolutionary mechanisms playing out over a complex biogeographic scenario in the Colombian Andes drove divergence in phenotypes and mitochondrial genomes of *Coeligena* hummingbirds, and lead to alternative hypotheses to be tested with whole-genome analyses.

## 1. Introduction

In the early days of sequence-based molecular systematics, mitochondrial DNA (mtDNA) was the marker of choice for most studies of population genetics, phylogenetics and phylogeography of animals because mtDNA is a haploid non-recombinant molecule almost free of non-coding regions, inherited via the maternal line, and abundant in tissues (Avise et al., 1987; Galtier et al., 2009; Wilson et al., 1985). Also, mtDNA evolves largely neutrally at a fast rate allowing one to find distinctive haplotypes among lineages (Avise et al., 1987; Ballard and Whitlock, 2004). However, mtDNA does not always reflect the evolutionary history of lineages owing to evolutionary and demographic processes such as selection, or differences between paternal and maternal dispersal and gene flow (Ballard and Melvin, 2010; Edwards et al., 2005; James et al., 2017). Thus, researchers have turned to assaying nuclear markers alongside mtDNA to study the divergence of lineages, an approach becoming increasingly feasible with the development of sequencing technologies allowing one to assay and analyze large numbers of genetic markers at relatively low cost (Kraus and Wink, 2015; Oyler-McCance et al., 2016; Toews et al., 2016). Information on genome-wide variation has not only contributed to more robust inferences of relationships among lineages as well as insights about how evolutionary mechanisms drive such divergence, but has also shed light on how evolutionary mechanisms interact to shape patterns of genetic divergence across genomes (Bonnet et al., 2017; Toews and Brelsford, 2012).

Phenotypes, nuclear genomes and mitochondrial genomes are not always equally divergent among lineages. When divergence is mainly driven by genetic drift, mtDNA is expected to diverge at a faster rate than nuclear DNA – and nuclear-encoded phenotypes – because the effective population size of the former is lower (Ballard and Whitlock, 2004; Moore, 1995). However, when selection drives divergence among populations, mtDNA need not diverge sooner than the nuclear genome, resulting in cases where patterns of mitochondrial and nuclear differentiation are not coincident or where phenotypic differentiation exists with little to no mitochondrial differentiation. Furthermore, phenotypically distinct populations may share mtDNA haplotypes because of mitochondrial introgression due to gene flow after divergence (Irwin et al., 2009; Rheindt et al., 2011; Toews and Brelsford, 2012).

Morphology, plumage, and songs are commonly used to compare populations and inform the species-level taxonomy of birds (Edwards et al., 2005; Remsen, 2005). Morphological measurements may provide evidence of barriers to gene flow (Cadena et al., 2018), whereas visual and acoustic signals are key phenotypes for species delimitation because they are involved in species recognition and reproductive isolation (Price, 2008; Roulin, 2004; Uy et al., 2009). Studies on Neotropical birds often show concordance in differentiation among lineages in phenotype and mitochondrial markers (e.g. Gutiérrez-Pinto et al., 2012; Lovette et al., 2010; Ribas et al., 2012; Sedano and Burns, 2010; Valderrama et al., 2014; Winger and Bates, 2015), although several examples exist of groups in which mtDNA is highly structured in distinct lineages despite little variation in plumage (Cadena et al., 2019; Chesser et al., 2020; D’Horta et al., 2013; Valderrama et al., 2014). Cases documenting species with marked differences in plumage coloration and little mitochondrial genetic divergence are more scarce (Campagna et al., 2012; Lougheed et al., 2013; Luna et al., 2017).

Among hummingbirds (Trochilidae), concordance between mtDNA divergence and overall differences in coloration between species and populations appears to be the norm (Chaves et al., 2007 *Adelomyia;* Jiménez and Ornelas, 2016 *Amazilia;* McGuire et al., 2008 Trochilidae; Ornelas et al., 2014 *Amazilia;* Parra et al., 2009 *Coeligena;* Zamudio-Beltrán and Hernández-Baños, 2018, 2015 *Laprolamia* and *Eugenes).* mtDNA divergence often coincides with differences in coloration among hummingbirds even when phenotypic variation is subtle, such as in the color of the crown, gorget, or tail (Benham and Witt, 2016 *Metallura;* Gonzalez et al., 2011 *C. curvipennis;* Lozano-Jaramillo et al., 2014 *Antocephala;* Ornelas et al., 2016 *Lampornis;* but see Rodríguez-Gómez and Ornelas, 2015 *Amazilia;* Sornoza-Molina et al., 2018 *Oreotrochilus).* There are, to our knowledge, only two documented cases of hummingbirds showing lack of genetic divergence with marked differentiation in coloration (i.e. differences in color in various plumage patches; Eliason et al., 2020; Parra, 2010), both occurring in the high Andes. One case involves two species of *Metallura* metaltails (Benham et al., 2015; García-Moreno et al., 1999, *Metallura theresiae* and *M. eupogon*) and the other two species of starfontlets in the genus *Coeligena* (Palacios et al., 2019; Parra et al., 2009) which we focus on in this study.

The Golden-bellied Starfrontlet (*C. bonapartei)* and the Blue-throated Starfrontlet (*C. helianthea)* inhabit the Northern Andes of Colombia and Venezuela (Figure 1A). The nominate subspecies of these species are sympatric in the southern part of their ranges in the Cordillera Oriental, whereas subspecies *C. b. consita* and *C. h. tamai* are allopatric in the Serranía de Perijá and Tamá Massif, respectively. These species are strikingly different in structural plumage coloration (Eliason et al., 2020; Sosa et al., 2020): *C. bonapartei* is greenish with fiery golden underparts whereas *C. helianthea* is blackish with a rose belly and aquamarine rump. Despite their markedly different phenotypes, *C. bonapartei* and *C. helianthea* are not genetically distinct in a mitochondrial gene (*ND2*), in a gene involved in the melanogenesis pathway (Melanocortin 1 Receptor *MC1R),* nor in regions flanking ultra-conserved elements (UCEs) across the nuclear genome (Palacios et al., 2019). Although these hummingbirds occupy similar environments, their lack of genetic differentiation is consistent with divergence with gene flow (Palacios et al., 2019). Phylogenetic analyses of sequences of the *ND2* mitochondrial gene also suggest that *C. b. consita* and *C. h. tamai* are more closely related to each other than either is to their nominate subspecies, a pattern more consistent with geography than with phenotype and taxonomy. However, it is unclear whether lack of genetic differentiation between *C. bonapartei* and *C. helianthea* is restricted to *ND2* or if it is a general pattern across the mitochondrial genome. Other mitochondrial markers may be more variable owing to differences among regions in substitution rates (e.g. *ND4* or the control region, Arcones et al., 2019; Eo and DeWoody, 2010) or in selective or stochastic demographic processes (Morales et al., 2015; Wort et al., 2017). Examining complete mitochondrial genomes might thus reveal heretofore undetected differences between species of *Coeligena*. Alternatively, if complete mitogenomes confirm lack of genetic divergence between *C. bonapartei* and *C. helianthea*, and that relationships of lineages of these species are inconsistent with their phenotype, then further consideration of mechanisms underlying evolutionary divergence in mtDNA and coloration in the group would be necessary. Such mechanisms potentially include natural and sexual selection as well as demographic processes acting during periods of geographic isolation and contact among lineages (Krosby and Rohwer, 2009; Morales et al., 2017; Pons et al., 2014; Toews et al., 2014).

**Figure 1.**
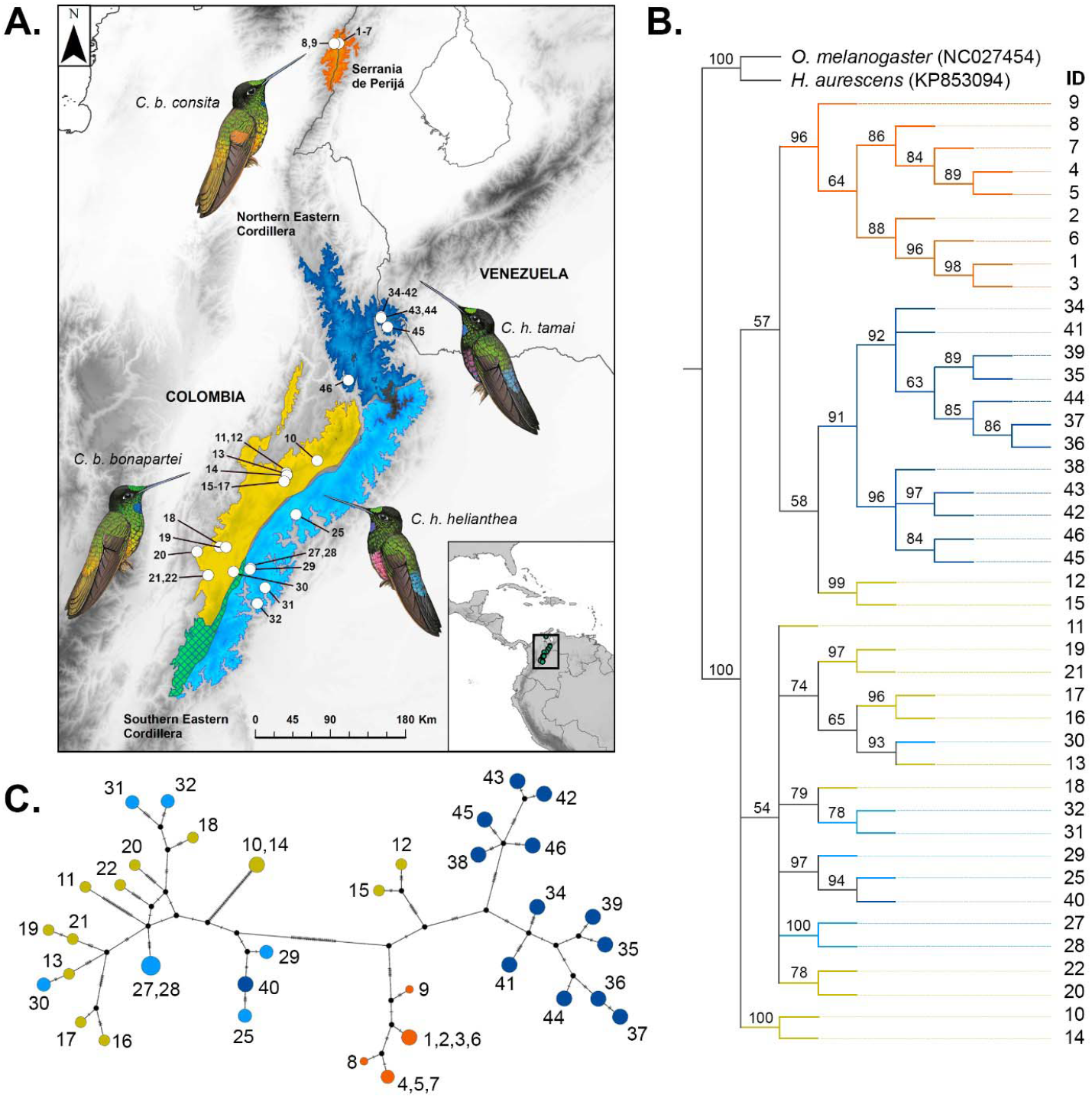
The maximum-likelihood phylogeny (B) and haplotype network (C) support two main mitogenome groups in *C. bonapartei* and *C. helianthea* more related to their geographical distribution (A) than with their taxonomic or phenotypic assignation. Note that the mitogenomes of *C. b. consita* and *C. h. tamai* are differentiated whereas the mitogenomes of *C. b. bonapartei* and *C. h helianthea* are indistinguishable. Numbers on the tips of the tree, on the haplotype network and on locations in the map correspond to individual IDs in Table S1. Colors correspond to the assigned subspecies *C. b. consita* (orange), *C. b. bonapartei* (yellow), *C. h. helianthea* (light blue), and *C. h. tamai* (dark blue). In the map the teal area is the region where nominate subspecies are sympatric. In the tree, numbers on branches are ML-bootstrap values; branch lengths were set to equal.

We sequenced and assembled complete mitochondrial genomes of multiple individuals to address the following questions: (1) Are the sequence and structure of the mitochondrial genomes of *C. bonapartei* and *C. helianthea* like those of mitogenomes of other bird and hummingbird species? (2) Is the lack of genetic divergence between *C. bonapartei* and *C. helianthea* a general pattern across the mitochondrial genome? (3) Are phylogenetic relationships of lineages of *C. bonapartei* and *C. helianthea* based on *ND2* also recovered using complete mitochondrial genomes? (4) Are different genes and regions in the mitochondrial genome equally informative about lineage relationships? And, (5) Are there substitutions in mitochondrial protein-coding genes among lineages of *C. bonapartei* and *C. helianthea* involving changes between aminoacids with different funcional characteristics which may suggest selection acting on these genes?

## 2. Material and methods

### 2.1. Samples and sequencing

We sampled 46 individuals, 23 each of *C. bonapartei* and *C. helianthea* (Supplementary Table 1), representing subspecies *C. b. bonapartei, C. b. consita, C. h. helianthea,* and *C. h. tamai*. Taxon identities were assigned by determination of specimens in the museum or by geography. Because previous work indicated that populations from the Mérida Cordillera of Venezuela often referred to *C. bonapartei* (subspecies *C. b. eos*) are genetically divergent from other populations in the complex (Palacios et al. 2019), we did not consider them in this study. Muscle tissue samples from voucher specimens were obtained from the collections of the Instituto Alexander von Humboldt (IAvH) and the Museo de Historia Natural de la Universidad de los Andes (ANDES). We employed relatively even samples sizes of each sex and subspecies of both *C. bonapartei* and *C. helianthea*.

We extracted total genomic DNA using a phenol/chloroform method and Phase-Lock Gel tubes, followed by a standard cleaning protocol employing magnetic beads. We prepared 46 Illumina TruSeq Nano DNA-enriched libraries following the manufacturer’s protocol for low-throughput configuration and 550bp insert size. We quantified the libraries using a Qubit fluorometer. Normalizing, pooling and sequencing were done by the Genomics Facility of the Institute of Biotechnology at Cornell University. Sequencing was performed using two lanes of NexSeq 500 2×150 paired end. We filtered the raw data by quality according to Illumina instructions, checked reads using Fastqc (Andrews, 2010), and cleaned them to remove adapters using AdapterRemoval (Schubert, Lindgreen, & Orlando, 2016).

### 2.2. Assembly and annotation of mitochondrial genomes

Although our sequence data contained sequences originating from both the nuclear and mitochondrial genomes, here we focus specifically on the later. We used MITObim v.1.9.1 (Hahn et al., 2013) with default parameters to assemble complete mitochondrial genomes from filtered reads following two alternative assembling strategies based on using different baits: (1) two independent assemblies using as baits the complete mitochondrial genomes of *Oreotrochilus melanogaster* and *Heliodoxa aurescens* (Genbank NC027454 and KP853094, respectively), and (2) a third assembly using as bait the *ND2* gene sequence for each individual -or a related one-available from previous work (Palacios et al., 2019). We expected that the first strategy would allow us to recover more complete individual mitogenome sequences because during initial iterations, reads would map to different sites on the reference mitogenome and this would allow extension from multiple edges. In turn, we expected that the gene-bait strategy would enable us to identify structural changes in genomes because it would allow extension only from the two edges of the gene, but it would likely be susceptible to recovering incomplete sequences when reads did not overlap, impeding continued extension.

The gene-bait strategy required multiple independent rounds of assembling. In each round we used as bait a new fragment obtained from the final genome assembled in the previous round. We compared the results from each strategy to determine the sequence and structure of mitogenomes of *C. bonapartei* and *C. helianthea*. In addition, we mapped the read-pool obtained from the complete-genome assembling strategy against the mitogenome sequence obtained from the gene-bait strategy using the “map to reference assemble” tool in Geneious 9.1.5 (http://www.geneious.com; Kearse et al., 2012). We used these map-to-reference assemblies to close gaps in some sequences, to check the number of repetitions at the end of the control region (see results), and to verify assigned alleles in each sequence at polymorphic sites. We aligned and edited mitochondrial genomes using ClustarO (Sievers et al., 2011) and manually in Geneious, and annotated them using MITOS beta version (http://mitos2.bioinf.uni-leipzig.de/index.py) and Geneious. In addition to the alignment of complete mitogenomes, for phylogenetic and population genetic analyses described below we generated alignments of each protein-coding gene (PCG), and a concatenated alignment of 13 PCGs *(ND1, ND2, COX1, COX2, ATP8, ATP6, COX3, ND3, ND4L, ND4, ND5, CYTB,* and *ND6*).

### 2.4. Population genetic, phylogenetic, and amino-acid change analyses

Using the alignment of complete mitogenomes, we calculated nucleotide diversity (Pi) for all sequences as a unit, and separately for *C. bonapartei, C. helianthea,* and for each of the four subspecies (*C. b. bonapartei, C. b. consita, C. h. helianthea, C. h. tamai*). We calculated absolute genetic divergence (Dxy) in DnaSP v6 (Rozas et al., 2017), and relative genetic divergence (Fst) between species and among subspecies assessing significance with 1,000 permutations using R package Hierfstat (Goudet and Jombart, 2015; R Core Team, 2017).

We examined phylogenetic relationships among individuals based on each of our alignments using maximum-likelihood analysis and computed majority-rule consensus trees in RAxML v8.2.12 (Stamatakis, 2014). We used the GTR+GAMMA model and multiparametric bootstrapping stopped by the autoMRE criterion. We used mitochondrial genomes of *Oreotrochilus melanogaster* and *Heliodoxa aurescens* (Genbank NC027454 and KP853094, respectively) as outgroups. We also built a median-joining haplotype network (Bandelt et al., 1999) in PopArt (Leigh and Bryant, 2015) using the complete mitogenome alignment.

Finally, we assessed whether there are fixed changes in amino-acids in proteins encoded in the mitogenome of lineages of *C. bonapartei* and *C. helianthea* potentially suggestive of selection. We first calculated the number and type of substitutions in each protein coding gene in DnaSP v6 (Rozas et al., 2017). Then, for each non-synonymous substitution we examined whether aminoacid variants were from different functional groups.

## 3. Results

### 3.1. Sequence and structure of mitochondrial genomes in C. bonapartei and C. helianthea

We recovered very similar sequence assemblies using the gene-bait and the complete mitogenome bait strategies. However, using the complete mitogenome strategy we observed insertions in some mitogenomes not recovered with the gene-bait strategy. Additionally, we found minor differences between assemblies obtained using the two strategies mainly in the length and sequence of the control region. We used the read-pool map-to-reference assemblies to resolve discrepancies between sequences from different assemblies and to review and manually correct nucleotide assignments in variant sites.

We recovered complete mitochondrial genomes for 42 of the 46 specimens (excluding IDs 23, 24, 26 and 33 in Supplementary Table 1, which we do not consider further because the data we obtained were of low quality), with an average coverage of 127.5x for all genomes (Max 1,555.7, Min 11.8, see Supplementary Table 1 for details, GenBank accession numbers XXX to XXX). The size of the mitochondrial genome of *C. bonapartei* and *C. helianthea* varied from 16,813 bp to 16,859 bp, mainly due to individual variation in length of a repetitive motive (‘AAAC’) at the end of the control region (beginning at 16,759 bp in the alignment). The 42 sequences were identical across 16,560 bp (98.2%), showed 248 variant sites (1.5%), with 51 positions having gaps or being ambiguous (0.3%). Mean pairwise identity was 99.7%, and total GC content was 44.8%. On average, the mitogenome sequences of *Coeligena* were identical to those of *O. melanogaster* across 14,555 bp (86.0%) and to those of *H. aurescens* across 14,450 bp (85.6%). The beginning of the control region (~350 bp) was the most difficult to align between sequences of *Coeligena* and those of outgroups. The mitochondrial genome structure of *Coeligena* species followed the typical pattern observed in other birds including hummingbirds, with 2 ribosomal RNAs, 13 protein coding genes, 22 transfer RNAs, and the control region (Figure 2).

**Figure 2.**
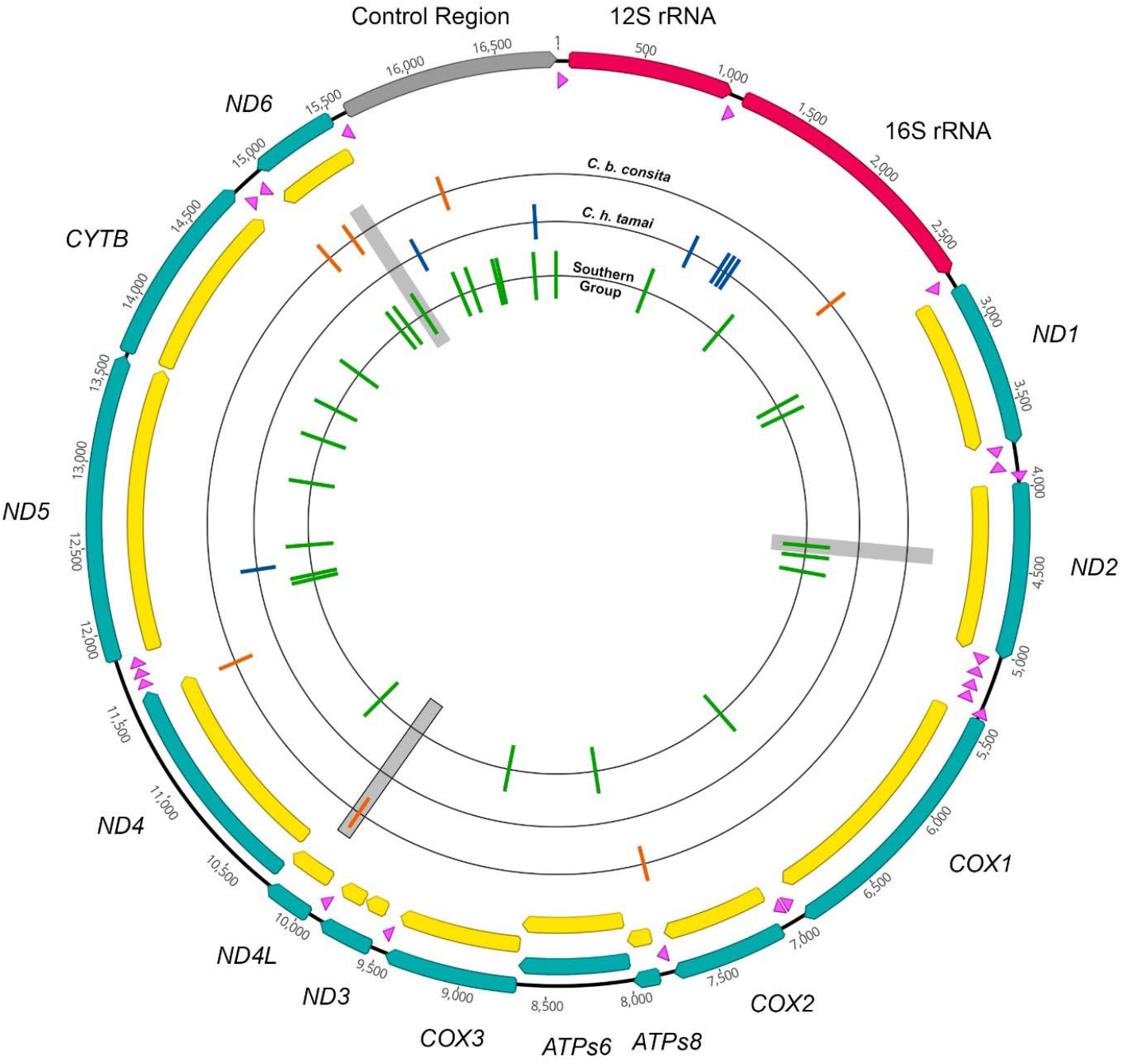
The mitochondrial genome structure of *Coeligena* hummingbirds follows the typical organization of birds: 22 tRNAS (pink), 2 rRNAs (raspberry), 13 protein-coding genes PCGs (blue), and the control region (gray). Coding sequences CDS are in yellow. Substitutions among the three genetic groups *C. b. consita* (orange), *C. b. tamai* (blue) and the southern group (green) are represented in the inner circles (singletons and intrapopulation variant sites are not represented). Gray boxes indicate the three non-synonymous substitutions found, the box with black edges indicates the only one involving a change between amino-acids with different functional features.

### 3.2. Genetic divergence and clustering patterns among lineages of C. bonapartei and C. helianthea

Across the complete mitogenome alignment including all individuals of *C. bonapartei* and *C. helianthea,* we found only 250 mutations (two sites had 3 alleles) in 248 variable sites (1.5% of the genome). Of these variable sites, 89 were singletons and 159 were parsimony-informative. Nucleotide diversity was low in the complete alignment (Pi = 0.00247, SD = 0.00013). The least diverse lineage was *C. b. consita* (Pi = 0.00019, 9 polymorphic sites), followed by *C. h. helianthea* (Pi = 0.00084, 40 polymorphic sites), *C. h. tamai* (Pi = 0.00124, 98 polymorphic sites), and *C. b. bonapartei* (Pi = 0.00254, 156 polymorphic sites). When we compared groupings based on species assignment (i.e. *C. bonapartei* vs *C. helianthea),* we found low relative genetic divergence (Fst = 0.076, p = 0.016). However, Fst values were greater when considering the four lineages separately (Table 1), with comparisons between lineages assigned to the same species showing higher relative genetic divergence than those between lineages assigned to different species (e.g. *C. b. consita* vs *C. b. bonapartei* Fst = 0.385, p-value < 0.001; *C. h. helianthea* vs *C. h. tamai* Fst = 0.518, p < 0.001; *C. b. bonapartei* vs *C. h. helianthea* Fst = 0.083, p = 0.1). All comparisons indicated low absolute genetic divergence (Dxy), supporting the general lack of genetic differentiation in the mitogenomes of these species (Table 1). However, high values of relative genetic divergence (Fst) between lineages of *C. bonapartei* and *C. helianthea* suggested genetic structure.

**Table 1.** Population genetic statistics and measures of genetic divergence between *C. bonapartei* and *C. helianthea* and among groups within. Nucleotide diversity Pi is lower in *C. b. consita* and *C. h. tamai* than in nominate subspecies. Absolute genetic divergence Dxy is low yet relative divergence Fst is high across comparisons. Genetic groups are derived from the clustering patterns analyses and are marked as “Gen” in the table.

| Population | # of Seq | # of Variants | Pi | Tajima's D | T's D p-value |
| --- | --- | --- | --- | --- | --- |
| *C. bonapartei* | 22 | 171 | 0.00249 | -0.44 | 0.351 |
| *C. helianthea* | 20 | 133 | 0.00218 | -0.09 | 0.481 |
| *C. b. consita* | 9 | 9 | 0.00019 | -0.05 | 0.500 |
| *C. b. bonapartei* | 13 | 156 | 0.00254 | -0.68 | 0.273 |
| *C. h. helianthea* | 7 | 40 | 0.00084 | -0.75 | 0.274 |
| *C. h. tamai* | 13 | 98 | 0.00124 | -1.54 | 0.057 |
| *C. b. consita* Gen | 9 | 9 | 0.00019 | -0.05 | 0.500 |
| *C. h. tamai* Gen | 12 | 54 | 0.00089 | -0.76 | 0.251 |
| Northern group Gen | 23 | 92 | 0.00120 | -0.76 | 0.242 |
| Southern group Gen | 17 | 100 | 0.00180 | -1.63 | 0.043 |

**Measures of genetic divergence**
| Population 1 | Population 2 | Fst | Fst p-value | Dxy |
| --- | --- | --- | --- | --- |
| *C. bonapartei* | *C. helianthea* | 0.076 | 0.0160 | 0.0026 |
| *C. b. consita* | *C. b. bonapartei* | 0.385 | 0.0010 | 0.0032 |
| *C. b. consita* | *C. h. helianthea* | 0.764 | 0.0010 | 0.0032 |
| *C. b. consita* | *C. h. tamai* | 0.402 | 0.0010 | 0.0017 |
| *C. b. bonapartei* | *C. h. helianthea* | 0.083 | 0.1069 | 0.0019 |
| *C. b. bonapartei* | *C. h. tamai* | 0.317 | 0.0010 | 0.0034 |
| *C. h. helianthea* | *C. h. tamai* | 0.518 | 0.0010 | 0.0033 |
| Northern group Gen | Southern group Gen | 0.514 | 0.0010 | 0.0035 |
| *C. b. consita* Gen | *C. h. tamai* Gen | 0.502 | 0.0010 | 0.0016 |

Phylogenetic analyses of the complete mitogenome alignment clustered all sequences of *Coeligena* hummingbirds in a well-supported clade (maximum-likelihood bootstrap ML-bs 100%, Figure 1). Relationships within this clade were unresolved, with a polytomy comprising (1) a clade grouping all sequences of *C. b. consita* (ML-bs 97%), (2) a clade grouping all but one of the sequences of *C. h. tamai* (ML-bs 90%), and (3) the remaining sequences (mostly of *C. b. bonapartei* and *C. h. helianthea*) scattered in smaller clades or by themselves. Phylogenies built with other alignments (each PCG and concatenated PCGs, Figure S1) showed lower resolution (i.e. more polytomies or lower support values). In most phylogenies, *C. b. consita* and *C. h. tamai* were more closely related to each other than either was to the nominate subspecies, but most support values for this grouping were lower than 80% except in the control-region phylogeny (ML-bs 88%).

All sequences of *C. b. consita* clustered together in the the median-joining haplotype network (Figure 1). All sequences but one of *C. h. tamai* clustered in another group which was close to, but distinguishable from, two sequences of *C. b. bonapartei* (ID 12 and 15). The remaining sequences of *C. b. bonapartei*, all sequences of *C. h. helianthea*, and the remaining sequence of *C. h. tamai* (ID 40) clustered in a third group (Figure 1). The network showed that sequences of *C. b. consita* and *C. h. tamai* are more similar to each other than to *C. b. bonapartei* and *C. h. helianthea*. Also, two individuals of *C. b. bonapartei* (ID 10 and 14) with the same haplotype were highly divergent from all other individuals. *C. b. consita* was the lineage with the lowest number of haplotypes (4 among 9 individuals). In the other lineages, the number of haplotypes was similar to the number of individuals: 12 haplotypes in *C. b. bonapartei* (13 individuals), 6 in *C. h. helianthea* (7 individuals), and 13 in *C. h. tamai* (13 individuals).

Based on the clustering patterns described above, we defined genetic groups for additional analyses in which we calculated the number of substitutions and measures of genetic divergence among groups. First, we defined (1) a northern group comprising all sequences of *C. b. consita*, all sequences of *C. h. tamai* except ID 40, and two sequences of *C. b. bonapartei* (ID 12 and 15); and (2) a southern group including most sequences of nominate subspecies *C. b. bonapartei* and *C. h. helianthea* (except ID 10 and 14) and one sequence of *C. h. tamai* (ID 40). Second, we considered separately the groups of *C. b consita* and *C. h. tamai* (excluding ID 40). There were only 27 substitutions (0.16%) yet high relative genetic divergence (Fst = 0.513, p-value < 0.001) between the northern and southern groups. Likewise, there were 14 substitutions (0.083%) and genetic divergence was high (Fst = 0.502, p-value < 0.001) between *C. b. consita* and *C. h. tamai.* The remaining 118 parsimony-informative sites existing among all sequences corresponded to intrapopulation diversity. Nucleotide diversity in the southern group (Pi = 0.0018) was higher than that of *C. b. consita* (Pi = 0.00019) and *C. h. tamai* (0.00089), but comparable to that of the northern group (Pi = 0.0012).

Given a substitution rate of 0.00256 substitutions per site per lineage per million years (s/s/l/My) for the complete mitogenome of birds (Eo and DeWoody, 2010), we estimated that the northern and southern groups diverged around 310,000 years ago, and that *C. b. consita* and *C. h. tamai* diverged around 160,000 years ago. Based on 13 protein-coding genes plus the two rRNAs and a substitution rate of 0.00164 s/s/l/My (mean rate for Apodiformes; Arcones et al., 2019) estimates of divergence times are similar yet slightly older: 380,000 years ago between the northern and southern groups, and 180,000 years ago between *C. b. consita* and *C. h. tamai*.

### 3.3. Functional aminoacid changes

Of the total 248 variant sites, 160 were located in protein-coding genes (Table S2). The remaining 88 variant sites were in rRNAs (6 in 12SrRNA, 20 in 16SrRNA), tRNAs (11), intergene spacers (5), and the control region (46). Among the 160 variant sites in protein-coding genes, 123 corresponded to synonymous changes and 38 to non-synonymous changes. Most non-synonymous changes were singletons (23 sites) or varied within populations (11 sites). Of the remaining 4 variant sites, a non-synonymous change was shared between one individual of *C. b. bonapartei* and one individual of *C. h. tamai* (T↔C position 269 in *ND5*). Only three non-synonymous changes corresponded to substitutions between genetic groups. One change in *ND2* and one in *ND6* were fixed differences between the northern and the southern groups (G↔A position 475 in *ND2,* and G↔A position 112 in *ND6).* These non-synonymous substitutions do not imply any evident functional changes because both aminoacids involved (valine and isoleucine) are aliphatic, nonpolar, and neutral. Finally, a non-synonymous substitution between *C. b. consita* and all other sequences (A↔G position 145 in *ND4)* implies a functional change in aminoacids. Whereas *C. b. bonapartei*, *C. h. helianthea* and *C. h. tamai* had the aliphatic, nonpolar alanine, *C. b. consita* had the hydroxyl-containing, polar threonine. Note that this change is not between the two main mitogenome groups because *C. h. tamai* has the variant of the southern mitogenenome group at this position.

## 4. Discussion

We found that the complete mitochondrial genomes of two hummingbird species differing strikingly in phenotype, *C. bonapartei* and *C. helianthea,* are highly similar. Mitogenomes of a sample of 42 individuals representing both species and two subspecies recognized within each of them were 98.2% identical. Moreover, estimates of genetic differentiation and clustering analyses of mitogenome sequences were unable to recover groups corresponding to species, and suggested instead that mitogenomes of *C. b. consita* and *C. h. tamai* formed distinct clusters more similar to each other than either was to mitogenomes of the nominate subspecies *C. b. bonapartei* and *C. h. helianthea* which were, in turn, indistinguishable from each other. These results indicate that patterns of variation based on the *ND2* gene (Palacios et al., 2019) are consistent across the mitochondrial genome, implying that the previously documented lack of mtDNA divergence between species does not reflect insufficient data nor atypical variation in *ND2* relative to other mitochondrial markers. Instead, patterns of variation and relationships among the mitochondrial genomes of the four lineages are inconsistent with phenotypic variation and current taxonomy, but seem to agree partly with geography, considering that *C. b. consita* and *C. h. tamai* occur in the Serranía de Perijá and the north of the Cordillera Oriental whereas both nominate subspecies occur to the south along the cordillera.

The discordance between mitochondrial genomes and coloration phenotypes in *C. bonapartei* and *C. helianthea* can be accounted for by various evolutionary processes which must have acted over a relatively short period of time given divergence-time estimates for the group. Based on the *ND2* gene, the clade formed by *C. bonapartei* and *C. helianthea* diverged from *C. b. eos* around 310,000 years ago, and the northern and southern clades comprising the four lineages of *C. bonapartei* and *C. helianthea* diverged around 240,000 years ago (Palacios et al., 2019). The latter estimate is more recent than our calculations of the divergence between the northern and southern groups at ca. 310,000 (complete mitogenome) or 380,000 years ago (PCG and rRNAs). Our estimates of divergence times must be interpreted with caution because different factors may bias them (Galtier et al., 2009; García-Moreno, 2004; Lovette, 2004), but they do suggest that the divergence between the northern and southern mitogenome groups, and the divergence between the mitogenomes of *C. b. consita* and *C. h. tamai* (160,000 estimated through complete mitogenomes and, 180,000 years ago using the PCG and rRNAs) are recent, i.e. happening within the past 500,000 years.

Ours is the first study in hummingbirds using complete mitochondrial genomes for a population-level analysis of genetic structure between species and across geography we are aware of, and few complete mitochondrial genomes of hummingbirds have been published (Morgan-Richards et al., 2008; Prosdocimi et al., 2016; Souto et al., 2016). We searched GenBank for complete mitochondrial genomes of closely related hummingbirds with more than a single individual sequenced per species to compare their divergence with the divergence we observed in *Coeligena*. We only found six mitogenome sequences for three subspecies of *Amazilia versicolor (A. v. versicolor* KF624601, NC_024156; *A. v. milleri* KP722042, NC033405; and *A. v. rondoniae* KP722041, NC_033404; Prosdocimi et al., 2016) representing populations occurring over a broad geographic range. Overall, these sequences are much more differentiated (5.1% of sites were variable) than our entire data set (1.5%). Although this comparison is far from comprehensive, it does support the idea that the mitogenomes of the lineages of *Coeligena* hummingbirds are highly similar and their divergence is quite recent relative to other hummingbirds with comparable data, as also indicated by analyses of individual mtDNA genes (Palacios et al., 2019; Parra et al., 2009).

In contrast to mtDNA phylogenies, nuclear markers suggest *C. b. consita* was the first branch to diverge in the group, whereas *C. h. helianthea* and *C. h. tamai* are reciprocally monophyletic groups forming a clade sister to *C. b. bonapartei* (Palacios et al., 2019; Palacios et al. unpublished). We found that complete mitogenomes of *C. b. bonapartei* and *C. h. helianthea* are undifferentiated even though both subspecies differ strikingly in phenotype and are also distinguishable using nuclear markers. Incomplete lineage sorting may explain this result because the southern mitogenome group exhibited high nucleotide diversity in comparison with *C. b. consita* and *C. h. tamai*, a pattern one would not expect due to a recent introgression (Krosby and Rohwer, 2009). However, nuclear sorting without mitochondrial sorting would be unlikely because the effective population size of the latter is ¼ that of the former. Instead, then, a scenario in which one mitogenome quickly swept through replacing the mitogenome of the other lineage and later recovered of nucleotide diversity may explain patterns of mitogenome sharing between *C. b. bonapartei* and *C. h. helianthea*.

The similarity in mitogenomes of *C. b. consita* and *C. h. tamai* appears more consistent with introgression after phenotypic differentiation in isolation. Mitochondrial introgression may often reflect selection (e.g. adaptive introgression via metabolic efficiency, Ballard and Melvin, 2010; Toews et al., 2014), but may also be due to demographic effects or to asymmetries between sexes in dispersal, mating behavior, and offspring production (Harris et al., 2018; James et al., 2016; Morales et al., 2017; Rheindt et al., 2014; Toews and Brelsford, 2012). We did not find functional changes in protein-coding genes between the northern and the southern mitogenomes suggesting adaptation, although adaptive changes related to substitutions in the control region (or in the 16SrRNA gen in the case of *C. h. tamai)* are possible. We are unaware of differential dispersal between sexes in *Coeligena*, in which dispersal and breeding biology are poorly known. Mitochondrial introgression between *C. b. consita* and *C. h. tamai* may have been facilitated by their geographical proximity and may have happened during a period of greater connectivity of forests in the Pleistocene (Flantua et al., 2019; Graham et al., 2010). Then, both lineages became isolated again and their mitogenomes diverged. The northern mitogenome may thus have evolved within *C. b. consita* and introgressed into *C. h. tamai* in a north to south direction, and such introgression may have further proceeded into *C. b. bonapartei* explaining why individuals ID 12 and 15 have haplotypes more closely related to the northern group.

The divergent mitogenomes of four individuals of *C. b. bonapartei* (ID 10, 12, 14, and 15) were unexpected considering the similarity among all other sequences. Although individuals ID 12 and 15 were closely related to the northern group, they shared 9 unique variants. Individuals ID10 and ID14 shared a mitogenome haplotype which was even more divergent (34 unique variants) sharing variants with both the northern (9) and the southern (18) groups. We can reject hybridization with other unstudied taxa as an explanation for these atypical mitogenomes because *ND2* sequences placed these specimens within the clade formed by *C. bonapartei* and *C. helianthea* to the exclusion of *C. b. eos* (Palacios et al. 2019). These atypical sequences may instead be evidence of persistence of a relict or a “ghost” mitochondrial lineage in *C. b. bonapartei* (Grandcolas et al., 2014; Zhang et al., 2019), which may have arisen and remained in isolation in the western slope of the Cordillera Oriental in Boyacá (Iguaque Massif and surroundings), a region where atypical patterns in mtDNA variation have been reported in other groups (Avendaño and Donegan, 2015; Chaves et al., 2011; Chaves and Smith, 2011; Chesser et al., 2020; Guarnizo et al., 2009). Another less likely explanation for these atypical sequences may be heteroplasmy and mitochondrial recombination which have been recognized in vertebrates in some cases (Piganeau et al., 2004; Rokas et al., 2003; Sammler et al., 2011).

In sum, based on our results and earlier work (Palacios et al. 2019) we hypothesize that a plausible evolutionary scenario accounting for patterns of mtDNA and phenotypic variation in *C. bonapartei* and *C. helianthea* is as follows. Based on comparison with the outgroup and other related species (*C. b. eos*, *C. lutetiae*, *C. orina*), the most probably body plumage coloration of the ancestor of our study clade was green with golden/orange underparts. The first lineage to diverge was likely *C. b. consita,* which evolved in the Serranía de Perijá in isolation from the ancestor of the other three lineages, retaining features of the ancestral plumage coloration but diverging in mtDNA. A second divergence event involved sister clades formed by *C. b. bonapartei* and *C. helianthea* (i.e. the common ancestor of both subspecies), with the former retaining the ancestral plumage and the latter evolving darker body coloration, rose belly, and aquamarine rump. These two lineages diverged in phenotype while maintaining an undifferentiated mitogenome owing to incomplete lineage sorting or introgression, except for populations of *C. b. bonapartei* which became isolated in the western slope of the Cordillera Oriental and diverged in mitogenome. Third, *C. h. tamai* and *C. h. helianthea* became isolated and diverged slightly in phenotype. Finally, during a period of forest connectivity the mitogenome of *C. b. consita* introgressed into *C. h. tamai,* a process followed by subsequent isolation of these lineages resulting in some divergence in their mitogenomes. Although this is a convoluted historical scenario, it is amenable to testing using genomic data and demographic models (e.g. Aguillon et al., 2018; Benham and Cheviron, 2019; Kearns et al., 2018) and other explanations for patterns of variation would appear even more complex.

## 5. Conclusion

Low genetic divergence among lineages of *C. bonapartei* and *C. helianthea* is a general pattern across their mitochondrial genomes despite their marked phenotypic differences. Mitogenomic variation in these lineages seems to more closely reflect geography and demographic history than the processes shaping their phenotypes and likely their nuclear genomes. Studying closely related lineages that diverged recently in complex topographic scenarios, such as the system of *C. bonapartei* and *C. helianthea*, might help to explain the different effects that evolutionary mechanisms may have in shaping the divergence between and within genomes. Incomplete lineage sorting, mitochondrial introgression, and demographic processes like population bottlenecks, phases of expansion and contraction, and the persistence of relict lineages have likely acted in this system resulting in marked discordance between phenotypes and mtDNA variation. A natural next step to understand the processes at work in this system is to place the results of the present study in the context of genome-wide patterns of genetic variation.

## Acknowledgments

We thank the Fundación para la Promoción del Banco de la República and the Lovette Lab at the Cornell Lab of Ornithology for financial support. For providing tissue samples we thank the Museo de Historia Natural de la Universidad de los Andes (ANDES) and the Instituto Alexander von Humboldt (IAvH). We exported tissues samples to the Cornell Lab of Ornithology (Ithaca, NY) thanks to CITES permit No. CO 41452 granted by the Ministerio de Ambiente y Desarrollo Sostenible of Colombia. We also thank Irby J. Lovette and Bronwyn G. Butcher for facilitating laboratory work. The manuscript was improved thanks to comments by XXXX. CP dedicates this paper to Tim Minchin.

## Supplementary material

**Figure S1.**
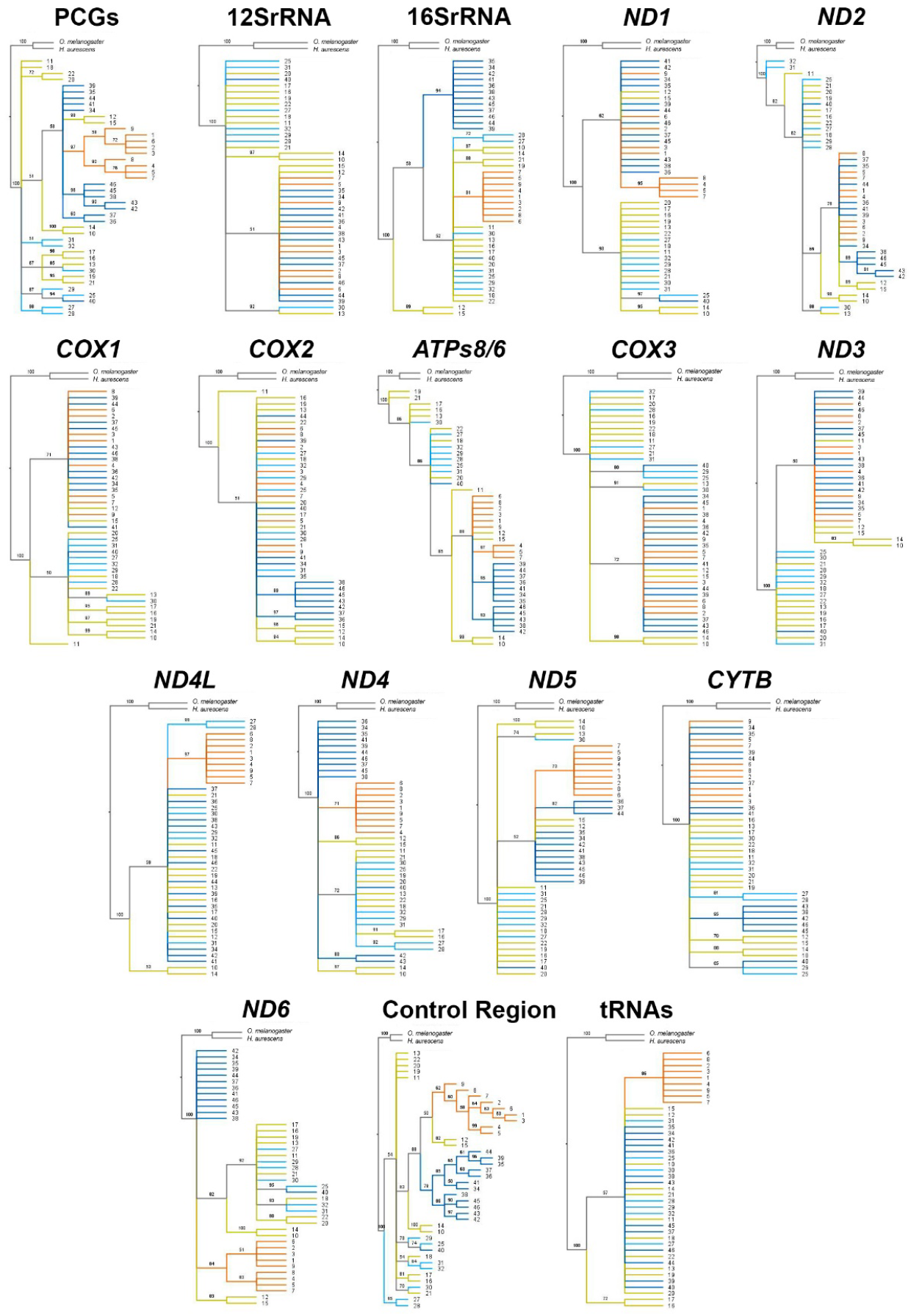
Maximun-likelihood phylogenies by alignment. Numbers on the tips of the trees correspond to individuals IDs in Table S1. Numbers on branches are ML-bootstrap values; branch lengths were set to equal in the trees. Colors correspond to the assigned subspecies *C. b. consita* (orange), *C. b. bonapartei* (yellow), *C. h. helianthea* (light blue), and *C. h. tamai* (dark blue).

**Table S1.**
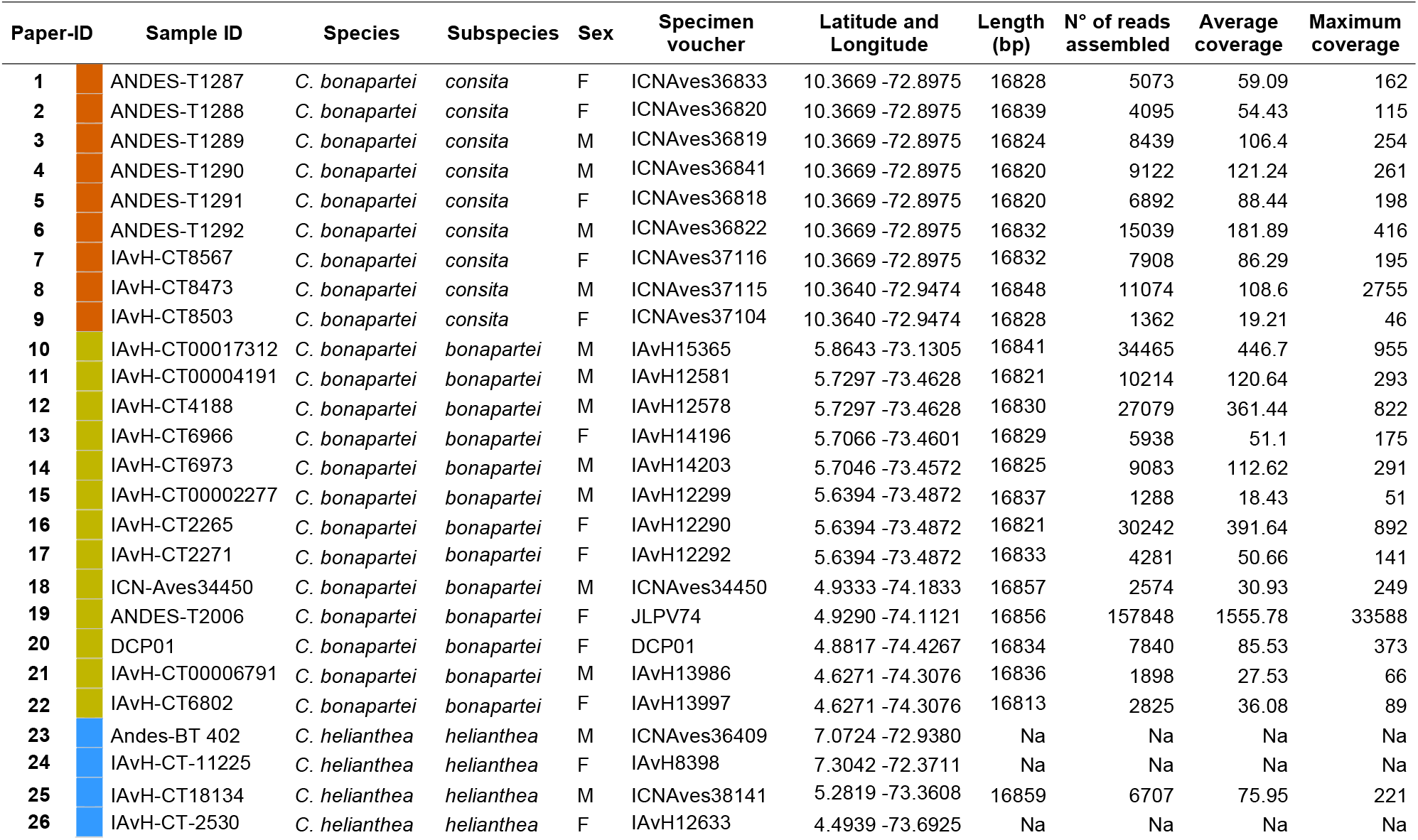
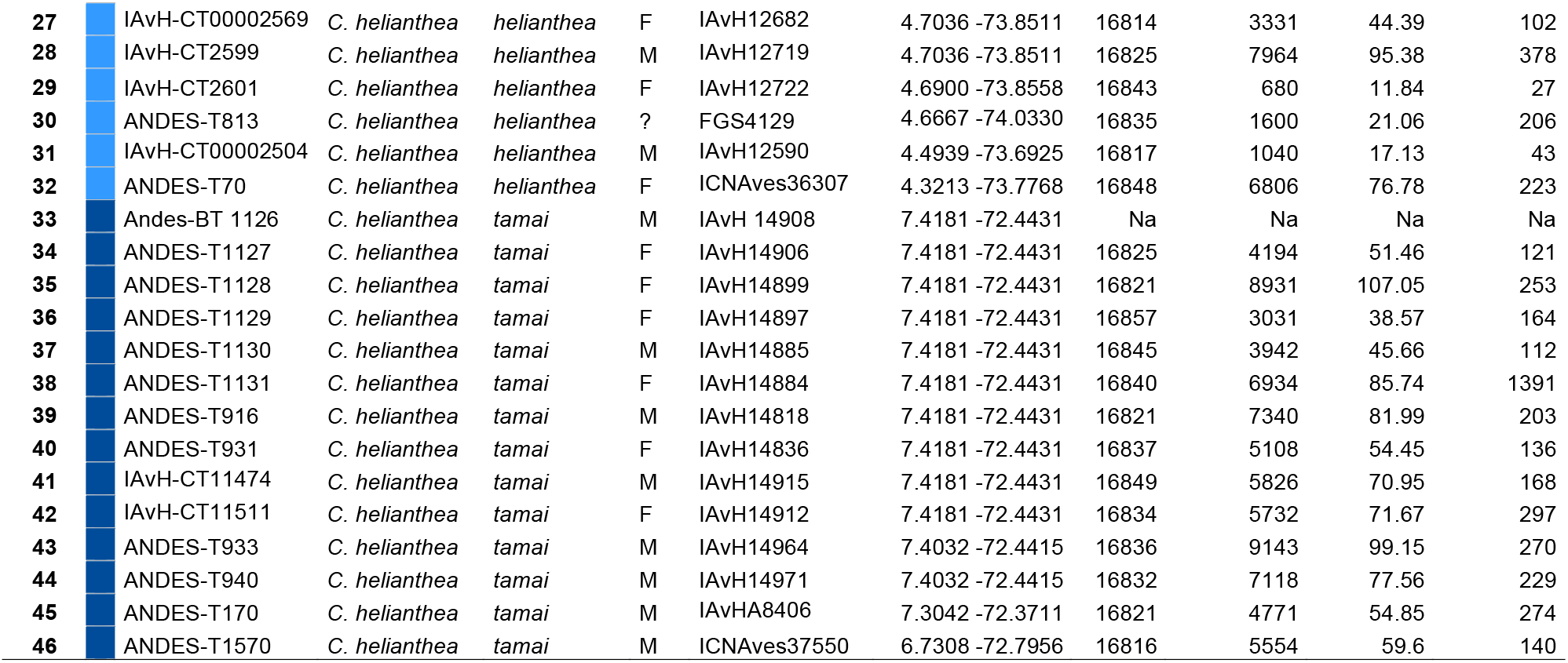
Specimen data and mitogenome assembly data. Paper-ID (identification number) corresponds to the ID used through this manuscript. Sample ID corresponds primarily to the tissue sample ID and secondarily to the specimen voucher or the collector’s ID. Specimen voucher column corresponds to the skin specimen ID in a collection or to the collector’s ID. All samples are from Colombia. Sex column F = female and M = male.

| Paper-ID | Sample ID | Species | Subspecies | Sex | Specimen voucher | Latitude and Longitude | Length (bp) | N° of reads assembled | Average coverage | Maximum coverage |
| --- | --- | --- | --- | --- | --- | --- | --- | --- | --- | --- |
| 1 | ANDES-T1287 | *C. bonapartei* | *consita* | F | ICNAves36833 | 10.3669 -72.8975 | 16828 | 5073 | 59.09 | 162 |
| 2 | ANDES-T1288 | *C. bonapartei* | *consita* | F | ICNAves36820 | 10.3669 -72.8975 | 16839 | 4095 | 54.43 | 115 |
| 3 | ANDES-T1289 | *C. bonapartei* | *consita* | M | ICNAves36819 | 10.3669 -72.8975 | 16824 | 8439 | 106.4 | 254 |
| 4 | ANDES-T1290 | *C. bonapartei* | *consita* | M | ICNAves36841 | 10.3669 -72.8975 | 16820 | 9122 | 121.24 | 261 |
| 5 | ANDES-T1291 | *C. bonapartei* | *consita* | F | ICNAves36818 | 10.3669 -72.8975 | 16820 | 6892 | 88.44 | 198 |
| 6 | ANDES-T1292 | *C. bonapartei* | *consita* | M | ICNAves36822 | 10.3669 -72.8975 | 16832 | 15039 | 181.89 | 416 |
| 7 | IAvH-CT8567 | *C. bonapartei* | *consita* | F | ICNAves37116 | 10.3669 -72.8975 | 16832 | 7908 | 86.29 | 195 |
| 8 | IAvH-CT8473 | *C. bonapartei* | *consita* | M | ICNAves37115 | 10.3640 -72.9474 | 16848 | 11074 | 108.6 | 2755 |
| 9 | IAvH-CT8503 | *C. bonapartei* | *consita* | F | ICNAves37104 | 10.3640 -72.9474 | 16828 | 1362 | 19.21 | 46 |
| 10 | IAvH-CT00017312 | *C. bonapartei* | *bonapartei* | M | IAvH15365 | 5.8643 -73.1305 | 16841 | 34465 | 446.7 | 955 |
| 11 | IAvH-CT00004191 | *C. bonapartei* | *bonapartei* | M | IAvH12581 | 5.7297 -73.4628 | 16821 | 10214 | 120.64 | 293 |
| 12 | IAvH-CT4188 | *C. bonapartei* | *bonapartei* | M | IAvH12578 | 5.7297 -73.4628 | 16830 | 27079 | 361.44 | 822 |
| 13 | IAvH-CT6966 | *C. bonapartei* | *bonapartei* | F | IAvH14196 | 5.7066 -73.4601 | 16829 | 5938 | 51.1 | 175 |
| 14 | IAvH-CT6973 | *C. bonapartei* | *bonapartei* | M | IAvH14203 | 5.7046 -73.4572 | 16825 | 9083 | 112.62 | 291 |
| 15 | IAvH-CT00002277 | *C. bonapartei* | *bonapartei* | M | IAvH12299 | 5.6394 -73.4872 | 16837 | 1288 | 18.43 | 51 |
| 16 | IAvH-CT2265 | *C. bonapartei* | *bonapartei* | F | IAvH12290 | 5.6394 -73.4872 | 16821 | 30242 | 391.64 | 892 |
| 17 | IAvH-CT2271 | *C. bonapartei* | *bonapartei* | F | IAvH12292 | 5.6394 -73.4872 | 16833 | 4281 | 50.66 | 141 |
| 18 | ICN-Aves34450 | *C. bonapartei* | *bonapartei* | M | ICNAves34450 | 4.9333 -74.1833 | 16857 | 2574 | 30.93 | 249 |
| 19 | ANDES-T2006 | *C. bonapartei* | *bonapartei* | F | JLPV74 | 4.9290 -74.1121 | 16856 | 157848 | 1555.78 | 33588 |
| 20 | DCP01 | *C. bonapartei* | *bonapartei* | F | DCP01 | 4.8817 -74.4267 | 16834 | 7840 | 85.53 | 373 |
| 21 | IAvH-CT00006791 | *C. bonapartei* | *bonapartei* | M | IAvH13986 | 4.6271 -74.3076 | 16836 | 1898 | 27.53 | 66 |
| 22 | IAvH-CT6802 | *C. bonapartei* | *bonapartei* | F | IAvH13997 | 4.6271 -74.3076 | 16813 | 2825 | 36.08 | 89 |
| 23 | Andes-BT 402 | *C. helianthea* | *helianthea* | M | ICNAves36409 | 7.0724 -72.9380 | Na | Na | Na | Na |
| 24 | IAvH-CT-11225 | *C. helianthea* | *helianthea* | F | IAvH8398 | 7.3042 -72.3711 | Na | Na | Na | Na |
| 25 | IAvH-CT18134 | *C. helianthea* | *helianthea* | M | ICNAves38141 | 5.2819 -73.3608 | 16859 | 6707 | 75.95 | 221 |
| 26 | IAvH-CT-2530 | *C. helianthea* | *helianthea* | F | IAvH12633 | 4.4939 -73.6925 | Na | Na | Na | Na |
| 27 | IAvH-CT00002569 | *C. helianthea* | *helianthea* | F | IAvH12682 | 4.7036 -73.8511 | 16814 | 3331 | 44.39 | 102 |
| 28 | IAvH-CT2599 | *C. helianthea* | *helianthea* | M | IAvH12719 | 4.7036 -73.8511 | 16825 | 7964 | 95.38 | 378 |
| 29 | IAvH-CT2601 | *C. helianthea* | *helianthea* | F | IAvH12722 | 4.6900 -73.8558 | 16843 | 680 | 11.84 | 27 |
| 30 | ANDES-T813 | *C. helianthea* | *helianthea* | ? | FGS4129 | 4.6667 -74.0330 | 16835 | 1600 | 21.06 | 206 |
| 31 | IAvH-CT00002504 | *C. helianthea* | *helianthea* | M | IAvH12590 | 4.4939 -73.6925 | 16817 | 1040 | 17.13 | 43 |
| 32 | ANDES-T70 | *C. helianthea* | *helianthea* | F | ICNAves36307 | 4.3213 -73.7768 | 16848 | 6806 | 76.78 | 223 |
| 33 | Andes-BT 1126 | *C. helianthea* | *tamai* | M | IAvH 14908 | 7.4181 -72.4431 | Na | Na | Na | Na |
| 34 | ANDES-T1127 | *C. helianthea* | *tamai* | F | IAvH14906 | 7.4181 -72.4431 | 16825 | 4194 | 51.46 | 121 |
| 35 | ANDES-T1128 | *C. helianthea* | *tamai* | F | IAvH14899 | 7.4181 -72.4431 | 16821 | 8931 | 107.05 | 253 |
| 36 | ANDES-T1129 | *C. helianthea* | *tamai* | F | IAvH14897 | 7.4181 -72.4431 | 16857 | 3031 | 38.57 | 164 |
| 37 | ANDES-T1130 | *C. helianthea* | *tamai* | M | IAvH14885 | 7.4181 -72.4431 | 16845 | 3942 | 45.66 | 112 |
| 38 | ANDES-T1131 | *C. helianthea* | *tamai* | F | IAvH14884 | 7.4181 -72.4431 | 16840 | 6934 | 85.74 | 1391 |
| 39 | ANDES-T916 | *C. helianthea* | *tamai* | M | IAvH14818 | 7.4181 -72.4431 | 16821 | 7340 | 81.99 | 203 |
| 40 | ANDES-T931 | *C. helianthea* | *tamai* | F | IAvH14836 | 7.4181 -72.4431 | 16837 | 5108 | 54.45 | 136 |
| 41 | IAvH-CT11474 | *C. helianthea* | *tamai* | M | IAvH14915 | 7.4181 -72.4431 | 16849 | 5826 | 70.95 | 168 |
| 42 | IAvH-CT11511 | *C. helianthea* | *tamai* | F | IAvH14912 | 7.4181 -72.4431 | 16834 | 5732 | 71.67 | 297 |
| 43 | ANDES-T933 | *C. helianthea* | *tamai* | M | IAvH14964 | 7.4032 -72.4415 | 16836 | 9143 | 99.15 | 270 |
| 44 | ANDES-T940 | *C. helianthea* | *tamai* | M | IAvH14971 | 7.4032 -72.4415 | 16832 | 7118 | 77.56 | 229 |
| 45 | ANDES-T170 | *C. helianthea* | *tamai* | M | IAvHA8406 | 7.3042 -72.3711 | 16821 | 4771 | 54.85 | 274 |
| 46 | ANDES-T1570 | *C. helianthea* | *tamai* | M | ICNAves37550 | 6.7308 -72.7956 | 16816 | 5554 | 59.6 | 140 |

**Table S2. Variant sites in the *C. bonapartei* and *C. helianthea* alignment. Excel file.**

